# CD36 family members are TCR-independent ligands for CD1 antigen-presenting molecules

**DOI:** 10.1101/2021.03.10.434884

**Authors:** Nicholas A. Gherardin, Samuel J. Redmond, Hamish E.G. McWilliam, Catarina F. Almeida, Katherine H.A. Gourley, Rebecca Seneviratna, Shihan Li, Robert De Rose, Catriona V. Nguyen-Robertson, Shian Su, Matthew E. Ritchie, Jose A. Villadangos, D. Branch Moody, Daniel G. Pellicci, Adam P. Uldrich, Dale I. Godfrey

**Author notes:** Correspondence: DIG, NAG. These authors contributed equally (Nicholas A. Gherardin, Dale I. Godfrey).

## Abstract

CD1c presents lipid-based antigens to CD1c-restricted T cells which are thought to be a major component of the human T cell pool. The study of CD1c-restricted T cells, however, is hampered by the presence of an abundantly expressed CD1c-binding partner on blood cells distinct to the T cell receptor (TCR), confounding analysis of TCR-mediated CD1c tetramer staining. Here, we identify the CD36 family (CD36, CD36-L1 and CD36-L2) as novel ligands for CD1c, CD1b and CD1d proteins, and show that CD36 is the receptor responsible for non-TCR-mediated CD1c tetramer staining of blood cells. Moreover, CD36-blockade enables tetramer-based identification of CD1c-restricted T cells and clarifies identification of CD1b- and CD1d-restricted T cells. We use this technique to characterise CD1c-restricted T cells ex vivo and show diverse phenotypic features, TCR repertoire and antigen-specific subsets. Accordingly, this work will enable further studies into the biology of CD1 and human CD1-restricted T cells.

**One Sentence Summary:** CD1 molecules bind CD36 family members and blockade of this interaction facilitates the study of CD1-restricted T cells.

## Introduction

T cell recognition of cognate antigen is a central event in adaptive immunity. Typically, the αβTCR, expressed on the surface of T cells, detects peptide-antigens presented by the polymorphic major histocompatibility complex (MHC) class I and II glycoproteins on the surface of antigen-presenting cells (APCs)^1^. This binding instigates rapid T cell activation and downstream effector function. An understanding of this receptor-ligand interaction led to the development of fluorescently labelled, recombinant peptide-MHC tetramer reagents that bind to T cells via their TCR^2^. This breakthrough permitted the tracking and isolation of Ag-specific T cell populations in human and mouse samples, thereby facilitating the rapid growth in our understanding of T cell biology, with ongoing implications in biomedicine.

Beyond peptide-antigens presented by MHC I and II, mammals have also evolved a series of monomorphic MHC I-like molecules that bind chemical moieties from other classes of antigenic material, including small cyclic metabolites presented by the MHC-related protein 1 (MR1), as well as lipids, presented by the CD1 family^3^. The human genome encodes genes for four distinct surface-expressed CD1 molecules including the group 1 CD1 genes *CD1A*, *CD1B*, and *CD1C,* and the group 2 CD1 gene *CD1D*, each of which encodes a protein with unique lipid-binding capacity^3, 4^. The four CD1 isoforms follow different intracellular trafficking routes, collectively sampling and presenting the cellular lipidome for T cell surveillance^5^. The amino acid diversity in the TCR binding site of each of the CD1 molecules engenders restriction by a discrete repertoire of T cells^3^. Accordingly, CD1a-, CD1b-, CD1c- and CD1d-restricted T cells play non-redundant roles in human health and disease^6^.

Most of our knowledge of CD1-restricted T cells pertains to a CD1d-restricted subset known as type I natural killer T (NKT) cells^7^. This is largely due to the high degree of evolutionary conservation of CD1d and the NKT TCR between mice and humans, along with the discovery of the archetypal CD1d-presented lipid-antigen α-galactosylceramide (α-GalCer)^8^ which is recognised by all NKT cell TCRs with high affinity^9^. This enabled the generation of α-GalCer-loaded CD1d tetramers that bind all NKT cells in humans and mice, and has facilitated their rapid characterisation^10, 11^, with mouse models highlighting key roles in diverse diseases including cancer, allergy and infection^12^. Conversely, group 1 CD1 proteins are not expressed in mice, and the TCR- and antigen-repertoires of group 1-CD1 restricted T cells appears to be more diverse^6^. Thus CD1a-, CD1b- and CD1c-restricted T cells remain much less extensively characterised. Most studies of group 1 CD1 have focussed on their role in the presentation of bacterial lipids including lipids from mycobacterium tuberculosis (Mtb)^13^. Indeed, mycobacterial-reactive CD8^+^ T cells from BCG-inoculated individuals are CD1-restricted^14^, and CD1 tetramers loaded with mycobacterial lipids have isolated CD1-lipid-specific T cells from human blood, including CD1a loaded with dideoxymycobactin (DDM)^15^, CD1b loaded with mycolic acid (MA)^16^ or glucose monomycolate (GMM)^17^ and CD1c loaded with phosphomycoketide (PM) and mannosyl phosphomycoketide (MPM)^18^.

An emerging paradigm suggests that group 1 CD1-self-lipid complexes are a major antigenic determinant for human T cells^19^. In some cases, CD1-restricted TCRs directly contact lipid antigens bound in CD1 clefts^20, 21^. More recent structural studies show that CD1a- and CD1c-autoreactive TCRs can directly bind the surface of CD1a^22, 23^ and CD1c^24^ when small ligands are buried within the CD1 molecule^22, 25, 26, 27, 28^. Indeed, CD1 tetramers loaded with the endogenous lipids from the mammalian cells used during their recombinant production (CD1-endo) have been employed to isolate CD1-autoreactive cells^22, 24, 28, 29^, and loading of CD1 molecules with specific self-lipids can modulate binding to CD1-autoreactive TCRs^20, 21, 22, 29^.

CD1c-autoreactive T cells have been suggested to be particularly abundant in human blood. Using limiting-dilution assays CD1c-autoreactive T cells accounted for up to 7% of CD4^+^ T cells in healthy adult donors^30^, and furthermore at least a subset of CD1-autoreactive T cells are detectable with CD1c-endo tetramers^24^. CD1c-reactive T cells also detect CD1c-expressing leukaemias, suggesting a role in surveillance of haematopoetic tumours^31^. Furthemore, CD1c is recognised by Vδ1^+^ γδ T cells^32^ as measured by clonal response assays and the use of CD1c tetramers^33^. Despite this progress, ex vivo detection of CD1c-restricted T cells is complicated by the fact that CD1c tetramers brightly stain some non-T cells in peripheral blood^24^. This finding suggests the existence of non-TCR ligands for CD1c and limits confidence in the identification and characterisation of CD1c-restricted T cells using tetramers. MHC and MHC-like molecules are known to have numerous binding partners, including killer cell immunoglobulin-like receptors (KIRs), natural killer receptors (NKRs), leukocyte immunoglobulin-like receptors (LILRs; also known as immunoglobulin-like transcripts; ILTs)^34^, and tetramer staining mediated by these molecules can hinder the enumeration and isolation of antigen-dependent T cell populations with tetramers. For example, HLA-E binds CD94-NKG2 receptors expressed by both NK and T cells^35^ and a monoclonal antibody (mAb) that blocks CD94/NKG2 receptors is required to abrogate this staining in order to isolate T cells that bind HLA-E via their TCRs^36^. Similarly, there is some evidence that CD1c and CD1d bind LILRB2 (ILT-4)^37, 38^, an immune receptor highly expressed on monocytes, however LILRB2 blockade only partially knocks down CD1 tetramer staining on monocytes suggesting that other ligands may also be present^37^.

Here, we demonstrate that CD1c binds to members of the CD36 family of lipid scavenger receptors including CD36 (SCARB3), CD36L1 (SCARB1) and CD36L2 (SCARB2/LIMP-2). Moreover, CD1b and CD1d also bind to these molecules albeit to a lesser extent. These findings reveal a previously unknown binding partner for CD1 family members in the human immune system that raises important new questions about the significance of this interaction. Furthermore, we show that this interaction is responsible for non-TCR mediated staining of human blood cells by CD1b, CD1c and CD1d tetramers, and that blockade of CD36 with anti-CD36 mAb allows the efficient identification of T cells expressing CD1b-, CD1c- and CD1d-reactive TCRs. This advance will open up new research into CD1-restricted T cells in general, and CD1c-restricted T cells in particular.

## Results

### Non-TCR-mediated CD1c tetramer staining is prevalent in human blood

To investigate CD1c-autoreactive T cells, we used a mammalian expression system to produce biotinylated, recombinant CD1 monomers which could then be tetramerised using fluorophore-conjugated streptavidin, as previously described^17^. These CD1 monomers furnish lipid-antigens derived from the cells used for expression and thus present a spectrum of endogenous mammalian lipids^39^. These tetramers are hence referred to as ‘CD1-endo’ tetramers and are used to detect ‘CD1-autoreactive’ cells. We generated BV421-conjugated CD1a-, CD1b-, CD1c-and CD1d-endo tetramers, as well as CD1d-α-GalCer and MR1-5-OP-RU tetramers that are used to detect NKT and MAIT cells, respectively^11, 40^. Each tetramer was used to stain human PBMC (**Fig 1A**). As expected^41^, MR1-5-OP-RU and CD1d-α-GalCer tetramers stained defined population of T cells. CD1a- and CD1b-endo tetramers provided no obvious staining relative to Streptavidin-BV421 alone. Confirming prior reports^24^, CD1c-endo tetramers, and to a lesser extent CD1d-endo tetramers, stained both CD3^+^ and CD3^-^ cells, suggesting that staining is not necessarily TCR-dependent.

**Figure 1:**
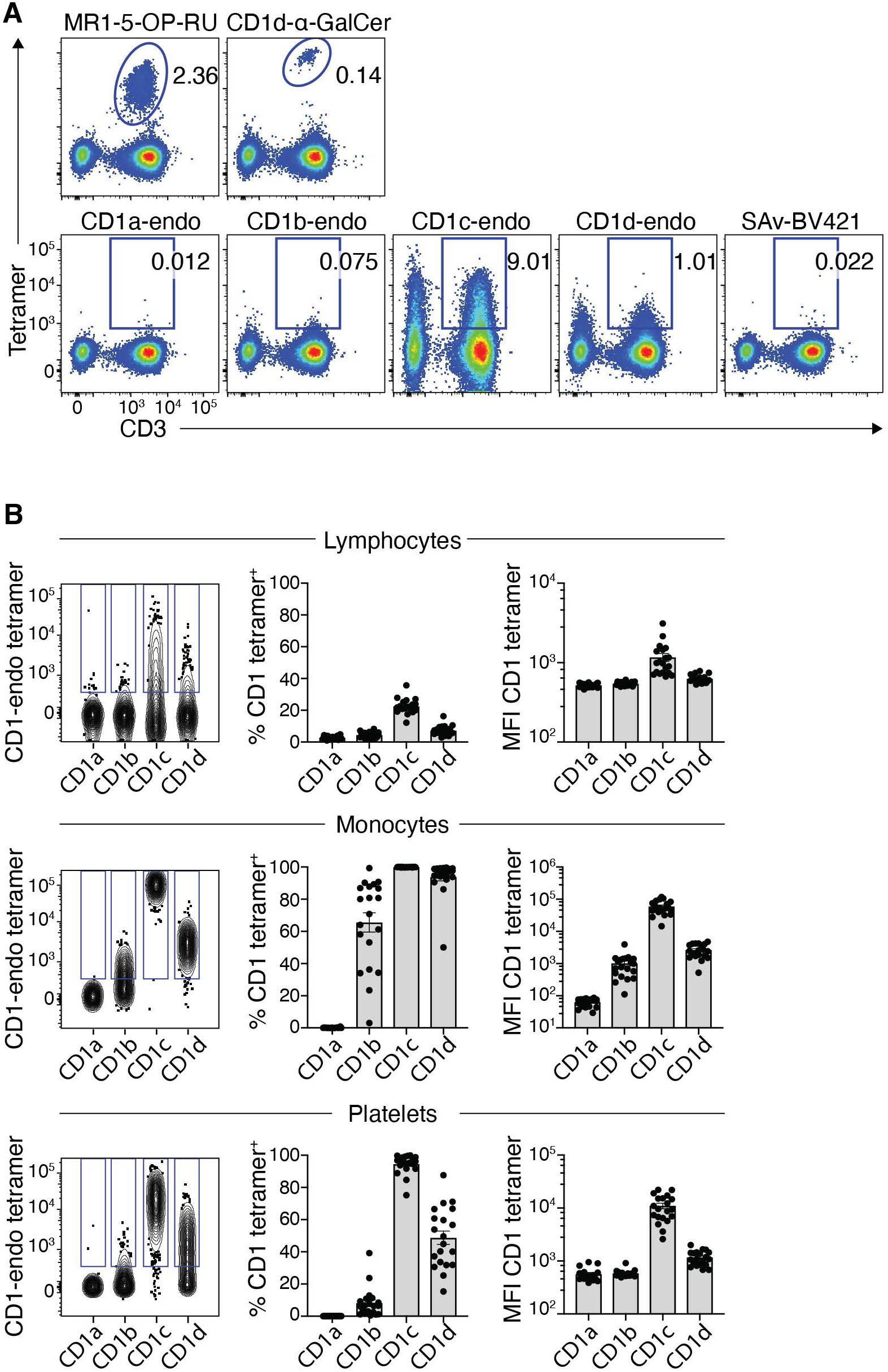
CD1-endo tetramers exhibit non-TCR-mediated binding to diverse blood cells. **A.** Example flow cytometric pseudo-colour plots showing tetramer staining on CD3^+^ T cells and CD3-lymphocytes using MR1-5-OP-RU, CD1d-αGalCer, CD1a-endo, CD1b-endo, CD1c-endo and CD1d-endo BV421 tetramers. **B.** Comparison of staining profiles of CD1-endo PE tetramers on lymphocytes (CD45^+^), monocytes (CD14^+^, SSC-A^hi^ and FSC-A^HI^) and platelets (CD42b^+^ SSC-A^low^ and FSC-A^low^). Left panel: contour plots showing concatenated CD1-endo tetramer staining profiles from the same donor. Different CD1-endo tetramers were used to stain separate samples and the data subsequently concatenated. Gates were set based on FMO. Middle panel: Bar graphs showing the proportion of cells staining with each CD1-endo tetramer (n=20). Right panel: Bar graphs showing median fluorescence intensity (MFI) of CD1-endo tetramer positive cells (n=20). SAv = streptavidin.

To further characterise the TCR-independent reactivity of CD1-endo tetramers, we stained human PBMCs and assessed the CD1-endo tetramer staining profile on distinct lineages of cells (**Fig. 1B**). CD1c, and to a lesser extent CD1d, exhibited a diffuse staining pattern, ranging from near background to very bright, on lymphoid cells. A median of 22% (range 12-36%) of lymphoid cells stained with CD1c-endo tetramers, and a median of 6.6% (range 3-13%) stained with CD1d-endo tetramers. Of those cells that stained with each tetramer, CD1c-endo tetramers were brighter relative to CD1d, with median median-fluorescence-intensities (MFI) of 988 and 613 respectively. Very little staining above background, as defined by fluorescence minus one (FMO), was detected with CD1a- and CD1b-endo tetramers, with medians of 2.6% and 4.6% respectively. Further analysis of NK, B and T lymphocyte subsets all gave similar staining patterns (**Fig. S1**). Monocytes were intensely reactive to CD1c-endo tetramers (MFI=52,000) and, to a lesser extent, CD1d-endo tetramers (MFI=2,431) (**Fig. 1B**). CD1b-endo tetramer staining was also apparent, albeit more variable, with a median of 75.4% (range 3-99%) and MFI of 866. In contrast, less than 1% of monocytes stained with CD1a-endo tetramers. Unexpectedly, platelets also reacted strongly with CD1b, CD1c and CD1d tetramers but not CD1a tetramers. These results suggest that CD1-endo tetramers bind to an undefined surface ligand that is widely expressed on cells of diverse lineages but is most abundant on monocytes and platelets, and to a lesser extent lymphocytes. Moreover, a consistent hierarchy of reactivity by the CD1 isoforms was apparent, with CD1c exhibiting the highest degree of reactivity, followed by CD1d, then CD1b, with very little evidence for CD1a-binding.

To further investigate T cell staining with CD1c tetramers, we reasoned that if the staining was TCR-mediated, we should be able to use flow cytometry to sort CD1c tetramer^+^ T cells and expanded them in vitro, yielding a population enriched for CD1c-endo tetramer^+^ T cells. However, after 10 days post-sort, most of the CD1c-endo tetramer staining was lost, despite these cells retaining CD3/TCR on their cell surface (**Fig. S2**). This supports an enrichment of non-TCR-mediated CD1c tetramer staining in ex vivo samples, and suggests that the ligand responsible for this staining is reduced with in vitro culture and/or expansion. This observation aligns with previous studies with CD1b ^20, 29, 42^ and CD1c^24^ tetramers that have successfully used this approach to isolate bone fide CD1-reactive cells from blood, although multiple rounds of sort-enrichment were required.

We next examninded the role of the α3 domain of CD1c in binding the undefined TCR-independet ligand(s), given that this domain is the key determinant in mediating MHC-binding to receptors such as LILRB2 (ILT-4)^43^. We therefore reasoned that by generating tetramers of chimeric CD1 proteins which have the α1 and α2 domains of CD1c fused to the α3 domain of CD1b, this will retain TCR CD1c-binding capacity, as previously published^44^, while heavily reducing any α3 domain-mediated, non-TCR binding. These chimeric tetramers stained Jurkat cells expressing a CD1c-restricted TCR but not a CD1b-restricted TCR (**Fig. S3A**), confirming that the chimeric protein indeed retained its specificity. However, the staining intensity on PBMCs was similar to that of non-chimeric CD1c tetramers (**Fig. S3B**), suggesting that non-TCR-mediated binding of CD1c to PBMC was dependent on the α1 and/or α2 domains.

### CD36 family members bind CD1 molecules

Given the range of primary cell types that stained with CD1c tetramers, we reasoned that some immortalised cell lines may also bind, providing a basis by which we could identify the candidate ligand(s). Indeed we found that C1R, 293T, K562, MEG-01 and THP-1 cells all stained uniformly and brightly with CD1c-endo tetramers, the brightest of which (K562) also stained weakly with CD1d-endo tetramers, suggesting that these cell lines are potential sources of CD1-binding ligand **Fig. 2A**.

**Figure 2:**
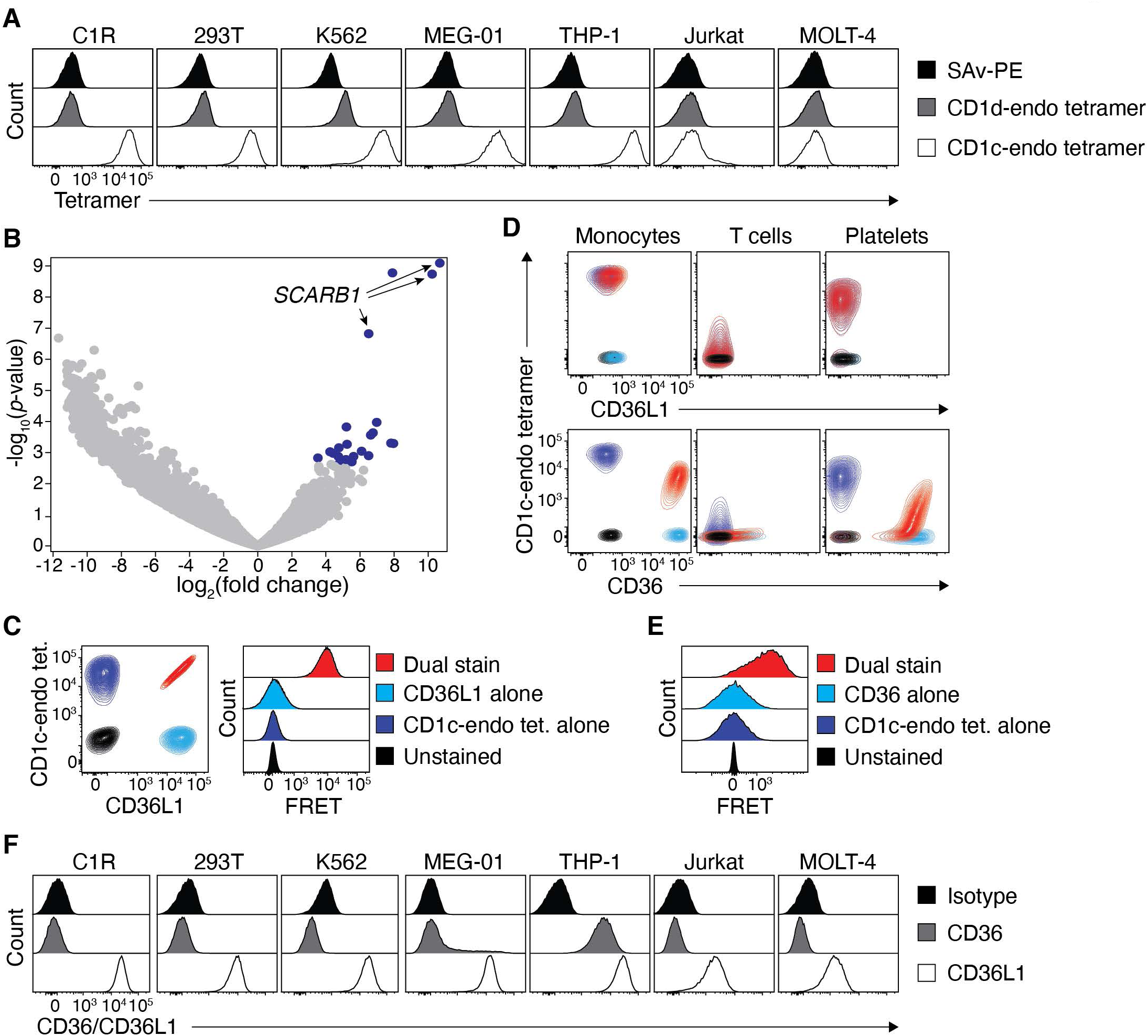
Identification of CD36 and CD36L1 as CD1 binding partners. **A.** Histogram overlays showing CD1c-and CD1d-endo tetramer staining on diverse cell lines. **B.** Volcano plot showing relative expression of gRNAs between C1R cells transduced with the GeCKO v1 human CRISPR knockout library versus those enriched for an inability to bind CD1c-endo tetramers. Blue data points represent guides significantly overrepresented with a false discovery rate <0.05 in the enriched pool. Arrows point to guides targeting the *SCARB1* gene. **C.** Left plot: Overlaid contour plots of C1R cells that were unstained (black), single labelled (navy/light blue) or co-labelled (red) with CD1c-endo tetramers and/or anti-CD36L1. Right plot: Histogram overlays of these cells showing emission in the FRET channel. **D.** Overlaid contour plots of blood cell subsets that were unstained (black), single labelled (navy/light blue) or co-labelled (red) with CD1c-endo tetramers and/or anti-CD36L1 (top panel) or anti-CD36 (bottom panel). Colour code as per C and E for CD36L1 and CD36 respectively (**E**) Histogram overlays of cells in (D) stained with anti-CD36, showing emission in the FRET channel. **F.** Histogram overlays showing CD36 and CD36L1 staining on diverse cell lines. All subfigures presenting FACS data are representative of 3 independent experiments.

To establish the identity of these CD1-binding ligand(s), we next performed a genome-wide CRISPR/Cas9-based library screen using C1R cells, which exhibited the highest reactivity to CD1c, and which have previously been used in a similar approach to identify genes that regulate surface expression of MR1^45^. After two rounds of FACS-purification of CD1c-endo tetramer^-^ C1R cells (ie. cells where the candidate ligand may have been deleted), this CD1c-endo tetramer^-^ population was enriched to ∼45% purity (**Fig. S4C**). Subsequent sequencing and analysis revealed 24 significantly enriched distinct guide RNA [false discovery rate (FDR) <0.05] (**Fig. S4D**). Of these, 3 of the top 4 guides targeted the *SCARB1* gene, with the remaining targeting *MBNL1*, a gene known to regulate alternative splicing of *SCARB1*^46^ (**Fig 2B**). The *SCARB1* gene encodes the CD36L1 protein (also known as SR-B1 and SCARB1), a surface-expressed member of the CD36 family of scavenger receptors involved in lipid metabolism with diverse roles in human physiology^47^. Given the lipid-binding capacity of CD36L1, this seemed a logical candidate binding partner for the lipid-presenting molecule CD1c. To test this hypothesis, we co-stained C1R cells with CD1c-endo tetramers and a mAb targeting CD36L1 (**Fig. 2C**, **left**). C1R cells stained brightly with anti-CD36L1 and the staining intensity was directly proportional to that of CD1c-endo tetramer. Moreover, when assessing Förster resonance energy transfer (FRET) between the donor fluorophore PE (conjugated to CD1c-endo tetramers) and acceptor fluorophore APC (conjugated to anti-CD36L1), a strong florescence signal was detected (**Fig. 2C**, **right**). These data suggest that CD1c-endo tetramers and anti-CD36L1 mAb are within 10 nm of each other on the cell surface. This supported the notion that CD36L1 is a direct ligand for CD1c and is responsible for CD1c-endo tetramer staining on C1R cells.

We next applied the same staining approach to PBMC, however, surprisingly, PBMC failed to stain with anti-CD36L1 (**Fig. 2D**, **top**). Since the CD36 family includes two other related receptors, CD36 (also known as SCARB3) and CD36L2 (also known as SCARB2 or LIMP2) we reasoned that these other CD36 family members may also facilitate binding to CD1. CD36L2 is not expressed on the cell surface and is restricted to the lysosome, however CD36 is known to be widely expressed on diverse cell types^48^. When staining PBMC with anti-CD36, we found that the staining profile on monocytes, T cells and platelets was highly similar to that of CD1c-endo tetramer staining (**Figure 2D**, **bottom**). Furthermore, when co-staining with anti-CD36 and CD1c tetramers, a clear coincident staining pattern was observed on both monocytes and platelets. Of note, in the presence of anti-CD36, CD1c-endo tetramer had reduced staining intensity, suggesting that the anti-CD36 mAb was partially blocking CD1c-endo from staining, which likely explains why the staining was only coincident on particularly brightly stained cells and was not observed on T cells. Akin to C1R cells, FRET was observed on monocytes costained with PE-conjugated CD1c-endo tetramer and APC-conjugated anti-CD36 mAb (**Fig. 2E**). Subsequently, we stained the original panel of cell lines for CD36 and CD36L1 expression and found that while all of the cell lines expressed CD36L1, the 2 lines that did not stain with CD1c-endo tetramers had approximately 10-fold less CD36L1 surface expression than the others (**Fig. 2F**). Only THP-1 cells also stained with anti-CD36. Collectively, these data suggest that CD36 expression on lymphocytes is responsible for non-TCR-mediated CD1c-endo tetramer staining on PBMC.

To confirm that CD36 family members were sufficient for binding to CD1 tetramers, we used a gene transfer approach whereby 293T cells were transiently transfected with plasmids encoding CD36 family members and then stained with a panel of CD1-endo tetramers. Since 293T cells already express CD36L1 (and stain with CD1c-endo tetramers) we first generated a 293T *SCARB1* knockout line using CRISPR/cas9 which sufficiently knocked-down CD1c-endo tetramer staining (**Fig. 3A**). These 293T.*SCARB1*^-/-^ cells were then transiently transfected to express full-length CD36, CD36L1 or CD36L2, and stained with CD1a, CD1b, CD1c and CD1d tetramers (**Fig. 3B**). Transfected cells specifically reacted with anti-CD36 family member mAbs, confirming their surface expression (**Fig. S5A**). Despite CD36L2 being a lysosomal protein, some residual protein was detected at the surface of CD36L2-transfected cells, which may reflect strong over-expression observed in this transfection system. Importantly, CD36 over-expressing cells stained clearly with CD1 tetramers, with a similar staining hierarchy to that observed with PBMCs, with CD1c-endo tetramers staining brightly, and CD1d- and CD1b-endo tetramers staining to a lesser degree (**Fig. 3B**). CD1a-endo tetramers failed to stain CD36-overexpressing cells. CD36L1 transfected cells also stained brightly with CD1c- and to a lesser extent CD1d-endo tetramers, whereas they failed to stain with CD1b and CD1a-endo tetramers. CD36L2-transfected cells only stained weakly with CD1c-endo tetramers, which may partly reflect its low surface expression. Control transfections of CD1-restricted auto-reactive TCRs were performed, and these showed TCR-dependent staining, thereby validating the integrity of the CD1-endo tetramers (**Fig. S5B**). Thus, all 3 CD36 family members can bind at least 1 CD1 family member.

**Figure 3:**
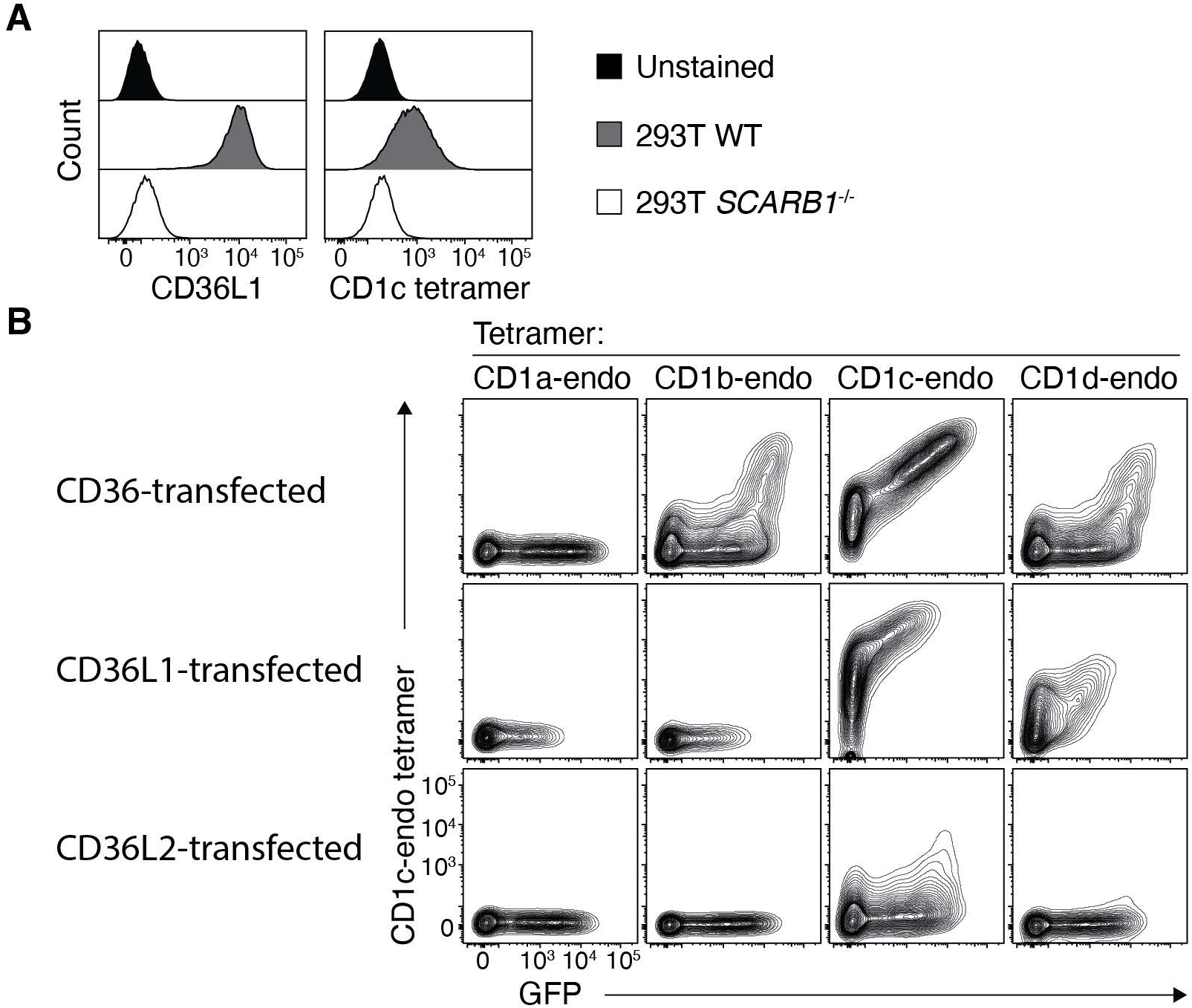
CD36 family members bind CD1 molecules. **A.** Histogram overlays showing CD36L1 and CD1c-endo tetramer staining on wildtype and *SCARB1*-deficient 293T cells. **B.** Contour plots showing CD1-endo tetramer staining on 293T.*SCARB1*^-/-^ cells transiently transfected to express CD36, CD36L1 or CD36L2. Data presented in (A) and (B) are representative of 3 independent experiments each.

### CD36 blockade facilitates characterisation of CD1c-restricted T cells

As observed above, co-staining with anti-CD36 mAb and CD1c-endo tetramers resulted in a reduction in staining intensity of non-TCR-mediated staining (**Fig. 2D**). These results suggested that the mAb and tetramers bind overlapping epitopes and that pre-incubation with anti-CD36 mAb may block CD36-mediated tetramer staining, leading to a clearer detection of T cell phenotypes that result from TCR-mediated CD1c binding. Indeed, titration of anti-CD36 pre-incubation resulted in a progressive reduction of CD1c-endo tetramer staining (**Fig. 4A-B**). A saturating dose of anti-CD36 blockade was observed at 3 μg/ml and this took the proportion of tetramer^+^ cells from a median of 40.5% when unblocked, to 0.02% when blocked (**Fig. 4A**). Thus, TCR-mediated tetramer staining accounts for a minor proportion of the staining observed in unblocked PBMC. This meant that after blocking, we should be able to explore the phenotypic features of fresh *ex vivo* CD1c-endo-restricted T cells. Given their rarity, we used magnetic-activated cell sorting (MACS) to enrich these cells from 6 healthy human PBMC samples (**Fig. 4C**). Consistent with CD1c being a prominent restriction element for γδ T cells^32, 33^, each donor yielded a mix of both αβ to γδ T cells. The rates of CD1c-endo tetramer staining for αβ versus γδ T cells was variable between donors, with a median of 17.9% γδ but range of 4.6-54.9% (**Fig 4D**). Focussing on the αβ lineage, the phenotypic features of the CD1c tetramer^+^ cells was diverse, spanning CD8^+^, CD4^+^ and double negative (DN) T cell subsets with a similar distribution to CD1c tetramer^-^ cells (**Fig. 4E**). Likewise, assessment of memory markers on the CD4 and CD8 subsets showed that CD1c tetramer^+^ cells were distributed similarly to tetramer^-^ cells across the 4 broad memory subsets found in blood (**Fig. 4F**). We next assessed the TCR-repertoire of these cells. Analysis of single cell-sorted CD1c-endo tetramer^+^ αβ T cells revealed diverse TRAV and TRBV gene usage with extensive diversity of CDR3α and CDR3β amino acid usage and length (**Table 1**). Despite this, we nonetheless observed an enrichment of TRBV4-1 in the TCR-β chain (6/26 matched pairs), in line with previous studies^49^ (**Fig. 4G**). We also noted an enrichment of TRAV17 (5/26) in the TCR-α chain, a variable gene known to be enriched in a population of CD1b-restricted T cells^50, 51^, as well as TRAV38-1 (4/26).

**Figure 4:**
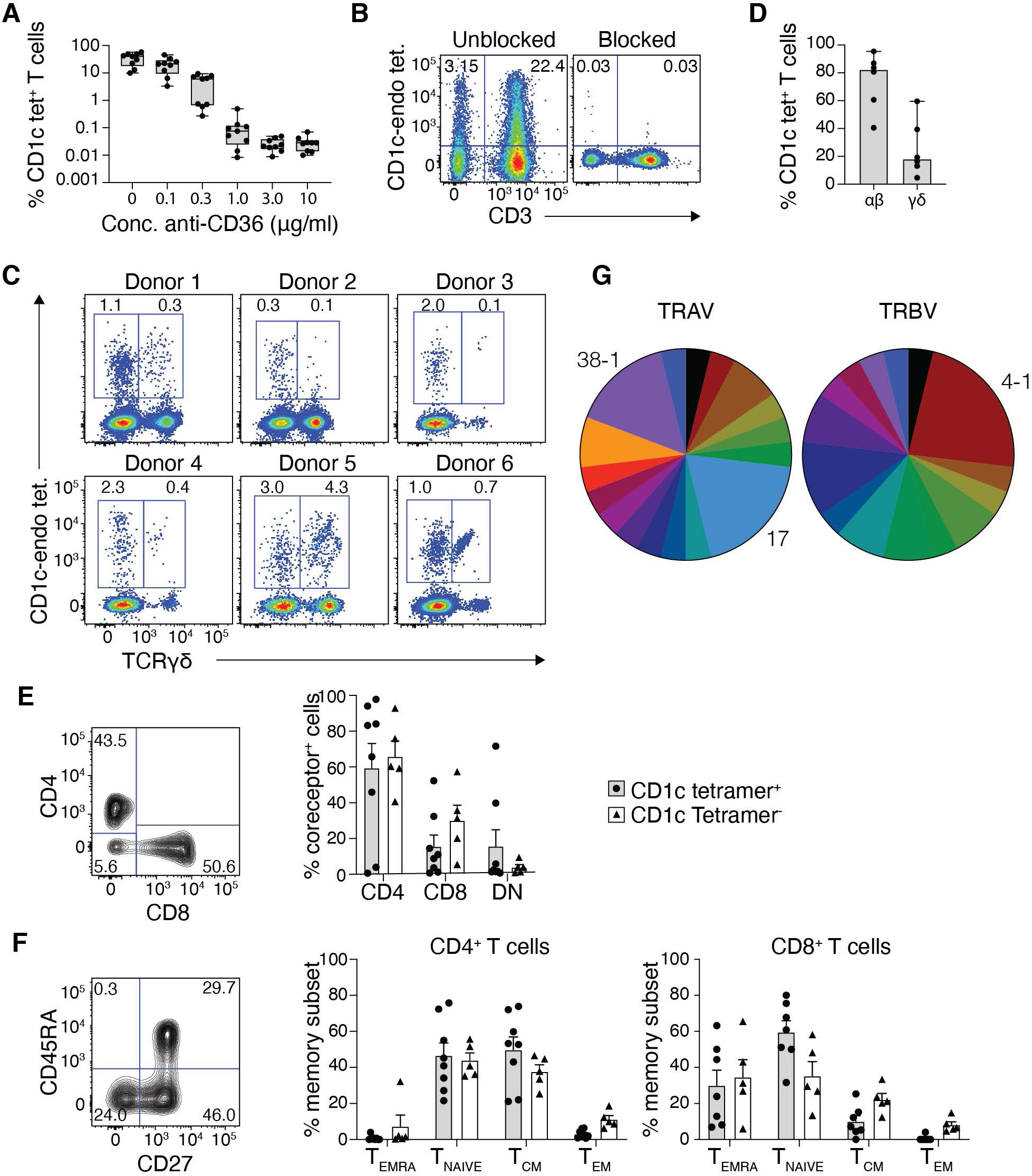
CD36 blockade enables characterisation of CD1c-reactive T cells. **A.** Bar graphs showing the proportion of CD3^+^ T cells in PBMC that stain with CD1c-endo tetramers after blocking with titrating doses of anti-CD36 (n=9). **B.** Example FACS plots showing CD1c-endo staining on lymphocytes with and without complete CD36 blockade. **C.** Example FACS plots gated on CD3^+^ T cells from 6 donor PBMC samples magnetically enriched with CD1c-endo tetramers. Plots show CD1c-endo tetramer staining on αβ (TCRγδ^-^) and γδ T cells. **D.** Bar graphs showing the proportion of magnetically enriched CD1c-endo-restricted T cells that utilise αβ or γδ TCRs (n=6 donors as shown in C). **E.** Coreceptor distribution of magnetically-enriched CD1c-endo-restricted αβ T cells. Left panel: representative FACS plot of CD4 and CD8 staining. Right panel: Bar graph showing coreceptor distribution from n=8 donors. **F.** Memory subset distribution of magnetically-enriched CD1c-endo-restricted αβ T cells. Left panel: representative FACS plot of CD27 and CD45RA staining. Middle and right panels: Bar graphs showing memory subset distribution of CD4 (middle) and CD8 (right) T cell subsets from n=8 donors. **G.** Pie charts showing distribution of distinct TRAV (left) and TRBV (right) genes used by n=26 CD1c-endo tetramer^+^ αβ T cells annotated in table 1. Enriched genes are labelled, remaining genes are differentiated by colour only.

**Table 1.**
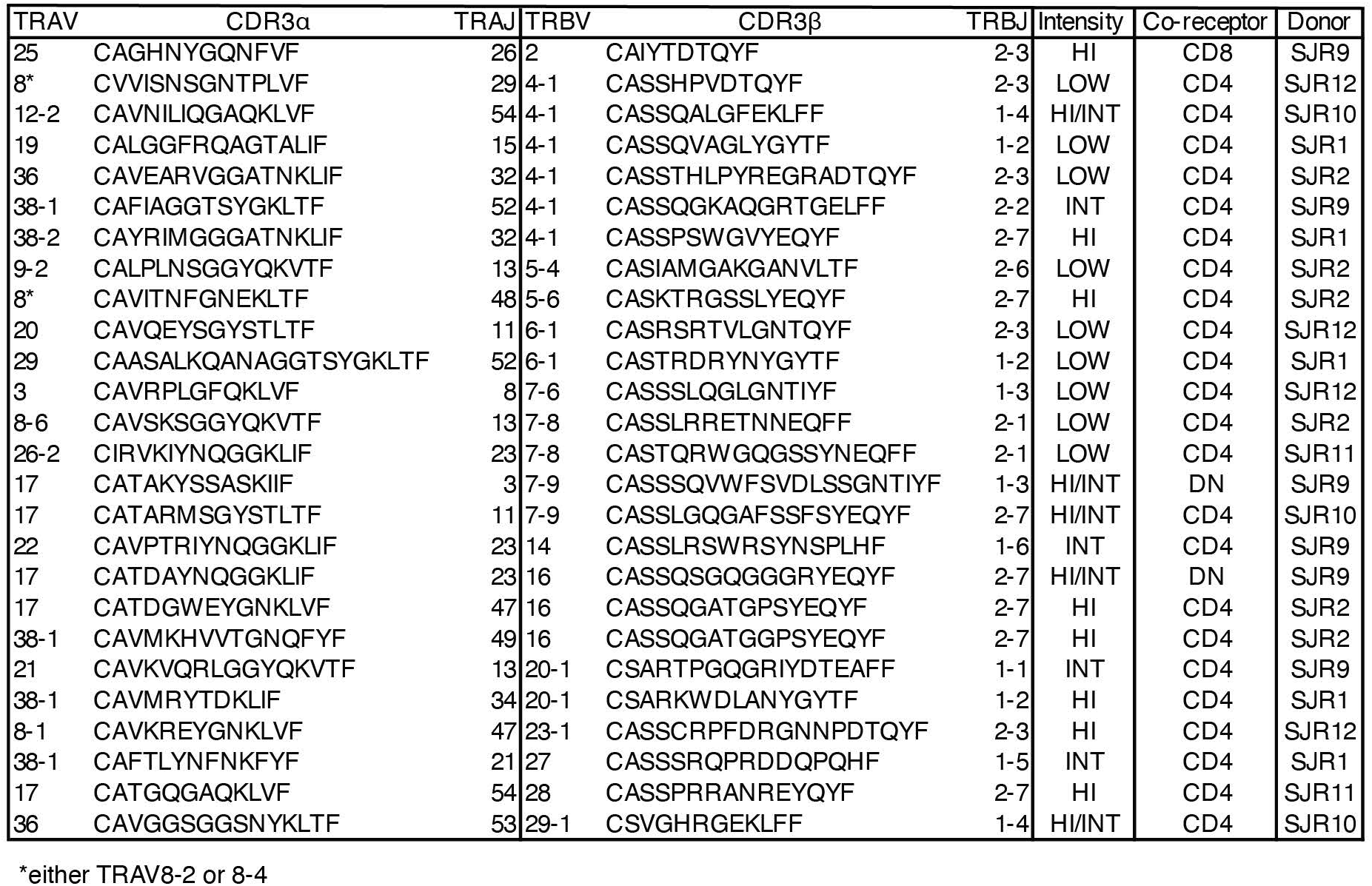
List of TCR sequences derived from CD1c-endo tetramer^+^ T cells

### Lipid-antigen discrimination by CD1c-autoreactive T cells

Having established a method of direct ex vivo staining, we next sought to determine whether we could isolate CD1c-autoreactive T cells with specificity toward specific self-lipids. We therefore loaded CD1c tetramers with lipids known to be presented by other CD1 isoforms, including the phospholipid, phosphatidylcholine (PC), the oxidised phospholipid, lysophosphatidylcholine (LPC), the sulfoglycolipid, sulfatide and GD3 ganglioside. Antigen-specific staining of T cells directly isolated from healthy blood was rare, but nonetheless, CD1c tetramer^+^ populations were observed with distinct staining hierarchies (**Fig 5A**). For example, donor 7 had a population of T cells that appeared to stain with all tetramers, whereas donor 8 had a population that was only present with CD1c-endo, vehicle, -PC and -LPC. On the other hand, donor 9 had the reciprocal staining pattern, with a population that preferentially stained with CD1c-sulfatide and -GD3. Different again, donor 10 predominantly stained with CD1c-sulfatide tetramers. To validate the antigen-specificity of these populations, we carried out single cell TCR analysis on samples of these CD1 tetramer^+^ T cells. While donors 8 and 10 yielded matched TCR-α and TCR-β chain sequences, both of which utilised TRBV4-1 (clones KG37 and KG52 respectively), donor 9 failed to yield a productive TCR-α chain. Noting that TRDV genes are frequently used by CD1d-restricted T cells (so-called δ/αβ cells)^52^, and CD1c teramers bind polycloncal Vδ1^+^ γδ T cells^33^, we repeated our sequencing incorporating the *TRDV* gene primers. This indeed yielded a productive *TRDV1* (encoding Vδ1) rearranged with the *TRAJ26* joining-gene and the *TRAC* TCR-α constant gene (clone NG30; **Fig 5B**). Transfer of these TCR genes into 293T.*SCARB1*^-/-^ cells and subsequently staining them with the same panel of CD1c tetramers recapitulated the lipid-reactivity observed in the primary cells (**Fig. 5C**). Thus, for example, NG30 derived from the CD1c-sulfatide and GD3-reactive cells in donor 9 was much more brightly stained by CD1c tetramers loaded with these versus other lipids. Although all CD1c tetramers stained the transfected cells to some extent, this likely reflects the high levels of TCR expressed in this model system. Collectively, these experiments not only validated the TCR-mediated CD1c-reactivity of CD1c tetramer^+^ cells after CD36 blockade, but also highlights a degree of self-lipid-antigen-discrimination in the CD1c-restricted T cell pool.

**Figure 5:**
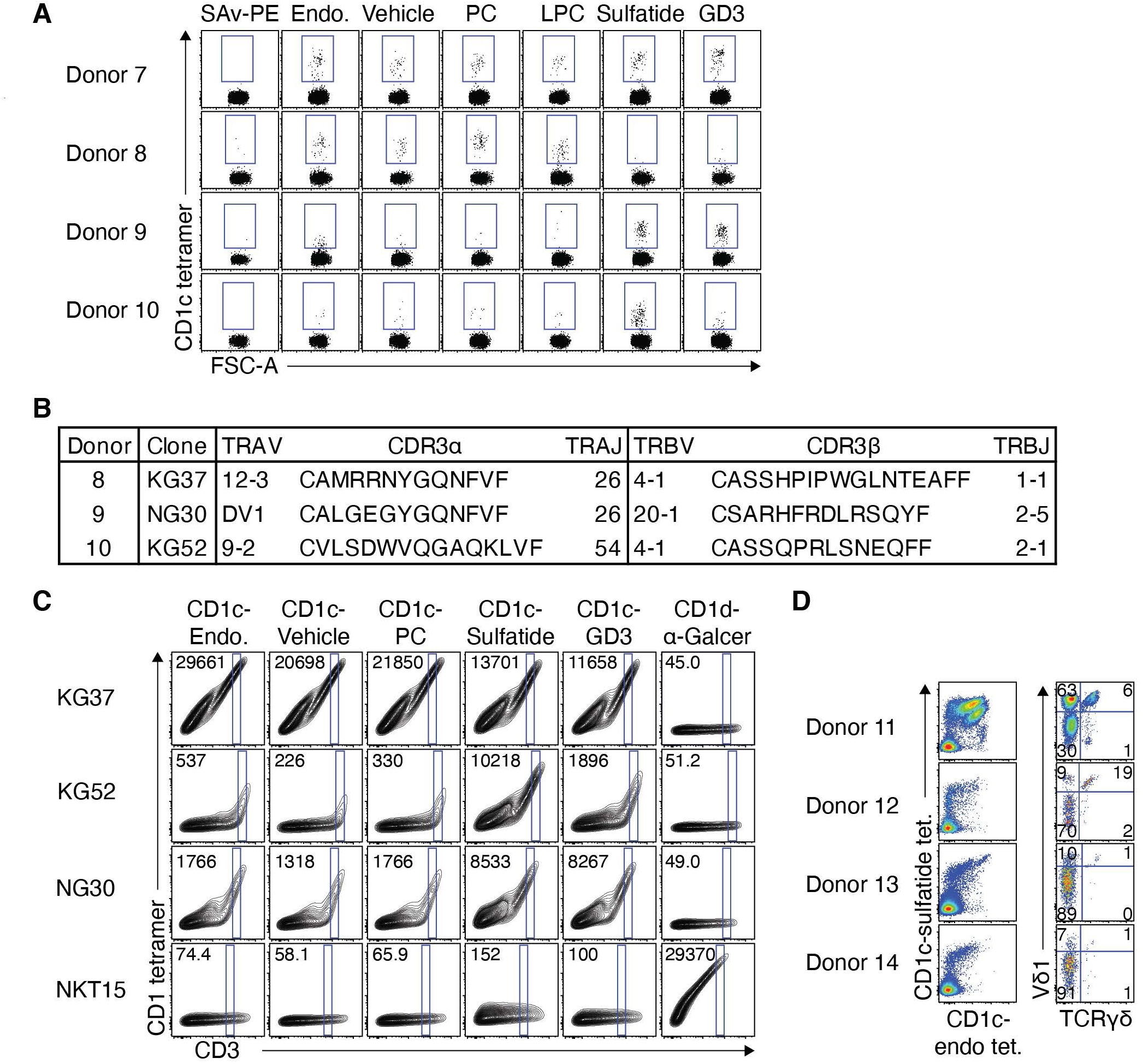
Lipid-antigen discrimination by CD1c-restricted T cells. **A.** Dot plots showing staining of CD3^+^ T cells from 4 donors with CD1c tetramers loaded with PC, LPC, sulfatide or GD3 compared to CD1c-endo or tyloxapol vehicle control. Streptavidin-PE (SAv-PE) was also used a negative staining control. **B.** Table showing TCR sequences derived from CD1c tetramer^+^ populations in figure A. **C.** Contour plots showing CD1-lipid tetramer staining of 293T.SCARB1^-/-^ cells transiently transfected to express CD1c-restricted TCRs from table in B or control NKT TCR clone NKT15. Consistent gates were set on a ‘window’ of CD3 expression across the different tetramer for each transfection, and median fluorescent intensity (MFI) of events in that gate is depicted in the top left of each plot. Experiment was performed 5 times for KG37, KG52 and NKT15 and 3 times for NG30, with similar results in each experiment. **D.** FACS plots from 4 donor PBMC samples co-stained and magnetically enriched with both CD1c-sulfatide and CD1c-endo tetramers. Left panel shows CD1c tetramer co-staining on enriched CD3^+^ T cells. Right panel shows TCRγδ and Vδ1 staining on total CD1c tetramer^+^ T cells.

Our finding that δ/αβ TCRs may be a frequent part of the CD1c-restricted T cell repertoire was intriguing. CD1c is a common restriction element for Vδ1^+^ γδ T cells^33^, and thus, Vδ1, whether in the context of a γδ or αβ TCR, may have a proclivity for CD1c, as it is known to have for CD1d^52^. To interrogate this finding further, we enriched T cells from 4 healthy PBMC samples, using both CD1c-sulfatide and CD1c-endo tetramers, and stained these cells with mAb against Vδ1 and pan-γδ TCR. When gating on T cells in the enriched pool (**Fig. 5D**, **left**), CD1c tetramer co-staining revealed distinct populations of CD1c-restricted T cells based on varying relative staining intensity with the 2 tetramers. For example, donor 11 had 2 populations of cells which stained brightly with both CD1c-endo and CD1c-sulfatide tetramers, but also some cells which preferentially bound CD1c-sulfatide or CD1c-endo tetramers. Donors 12, 13 and 14 lacked a prominent population of CD1c-endo exclusivity but had populations that appeared to preference CD1c-sulfatide or co-stained with both tetramers. When gating on total CD1c tetramer^+^ cells and assessing the TCR usage (**Fig. 5D**, **right**), the distribution of subsets was variable between the 4 donors. γδ T cells accounted for 7%, 21%, 2% and 2% for donors 11-14 respectively, and Vδ1^+^ T cells accounted for 69%, 27%, 11% and 8%. When focussing on Vδ1^+^ δ/αβ T cells, these accounted for 63%, 9%, 10% and 7% of total CD1c tetramer^+^ cells, and 91%, 47%, 91% and 88% of the Vδ1^+^ fraction for donors 11-14 respectively. Accordingly, like CD1d-restricted T cells^53, 54, 55^, Vδ1 is frequently used in both the αβ and γδ CD1c-restricted T cell repertoires.

### CD36 blockade facilitates isolation of CD1d- and CD1b-restricted T cells

Given CD1d-endo and CD1b-endo tetramers also bind PBMC in a CD36-dependent manner, albeit to a lesser degree than CD1c-endo tetramers, we reasoned that CD36 blockade may also be a valuable tool for the *ex vivo* isolation of CD1d-restricted type II NKT cells and CD1b-restricted T cells including GMM-reactive germline-encoded mycolyl-lipid-reactive (GEM) T cells^56^. To test this, we first stained healthy PBMC from 3 donors with CD1d-endo, CD1d-LPC and CD1d-sulfatide tetramers, using CD1d-α-GalCer tetramers as a control (**Fig. 6A**). In line with the staining profile in figure 1, the identity of the loaded-lipid affected the level of CD36-mediated staining; CD1d-endo and CD1d-LPC tetramers had substantial CD36-mediated staining on both T and non-T cells, whereas this was less prominent in the CD1d tetramers loaded with sulfatide or α-GalCer, suggesting that some bound lipids may inhibit CD36 binding to CD1d. Nonetheless, CD36-blockade prior to tetramer staining resulted in a marked reduction in staining events, revealing a rare residual population of tetramer^+^ T cells in each case. Thus, while CD36-blockade may be less necessary for staining of NKT cells with CD1d-α-GalCer, CD36-blockade may facilitate the ex vivo analysis of CD1d-restricted cells with type II Ags such as sulfatide and LPC^57^. We next took a similar approach, but instead used CD1b-GMM tetramers (**Fig 6B**). While CD1b tetramers did not display the same level of non-TCR-mediated staining as did CD1c and CD1d tetramers, there was nonetheless a notable improvement in signal:noise for analysis of TRAV1-2^+^ GEM T cells. For example, donors 18 and 20 had prominent populations of GEM T cells which were more readily defined after CD36-blockade. Reciprocally, the rare TRAV1-2^+^ cells that stained with CD1b-GMM tetramers in donor 19 where no longer present after CD36-blockade. Accordingly, CD36-blockade is also applicable to analysis of CD1b-restricted T cells, particularly rare populations where high signal:noise is required.

**Figure 6:**
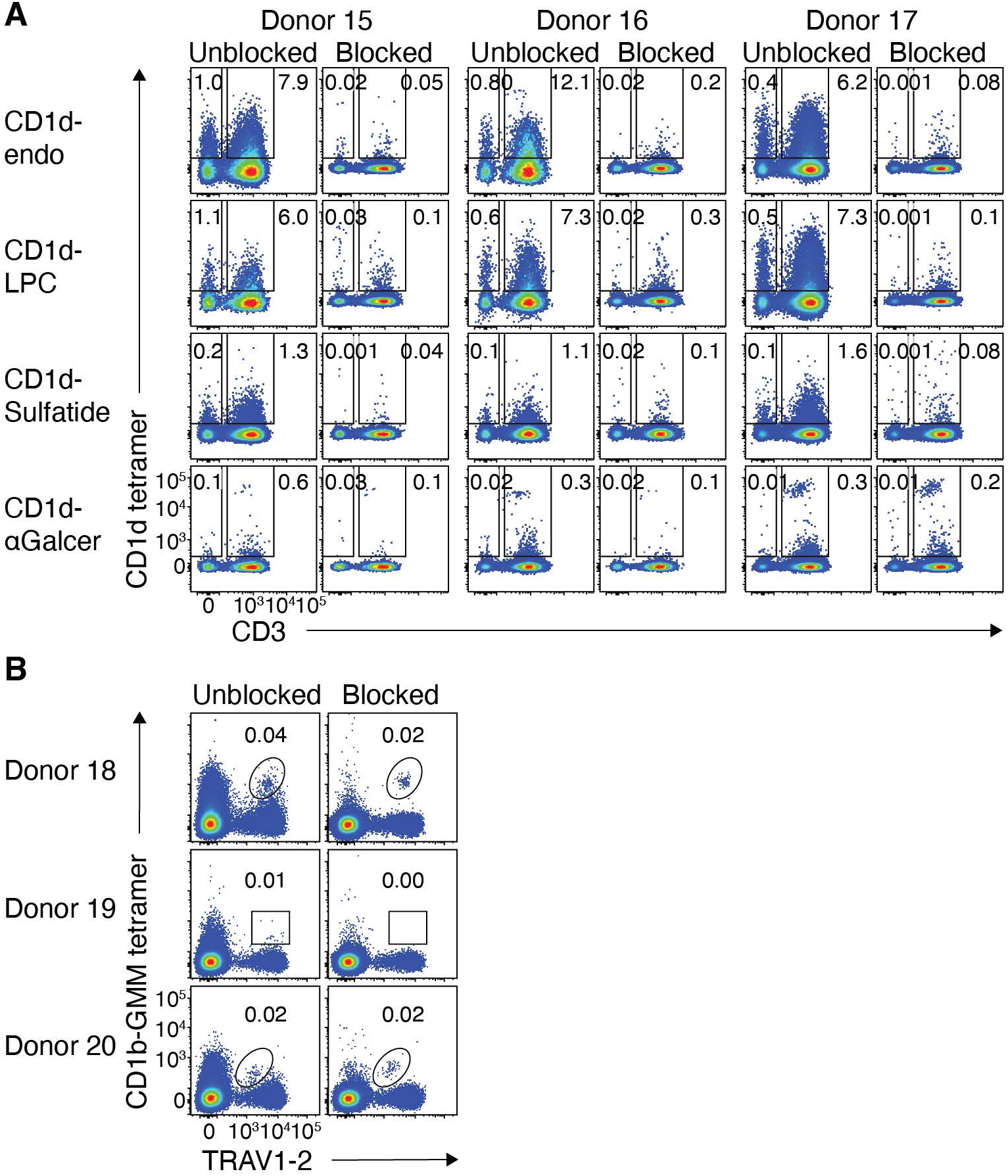
CD36-blockade facilitates ex vivo analysis of type II NKT cells and GEM-T cells. **A.** FACS plots from 3 healthy donor PBMC samples (representative of a cohort of 8) showing CD1d tetramer staining on lymphocytes with and without CD36-blockade. **B.** FACS plots from 3 healthy donor PBMC samples showing CD1b-GMM tetramer staining on CD4^+^ T cells with and without CD36-blockade.

## Discussion

The primary function of MHC and MHC-like molecules, such as CD1 family members, is thought to be the presentation of antigens to TCRs expressed by T cells^1^. However, these surface proteins can also interact with other non-TCR molecules, such as KIR and LILR family members, which appears to be important in modulating immune responses and preventing subversion of immune responses by unstable tumours and MHC-modulating viruses^34^. Here, we have identified previously unrecognised interactions between two important families of surface receptors, being the CD1 and CD36 molecules. While the functional significance of this interaction remains to be explored, an obvious association is the lipid binding role of both families of molecules. The CD36 family regulates lipid metabolism through binding of cholesterols, fatty acids and normal and modified lipoproteins^47^. It is thus tempting to speculate that the CD36 family may have a role in lipid-Ag-presentation by CD1 molecules, a process known to involve other lipid scavenger receptors^58^. Indeed, apolipoproteins, a component of lipoproteins that are scavenged by CD36, have been shown to be important in the efficient delivery of exogenous lipids-Ags to APCs to mediate NKT cell activation^59^. Thus, CD36 family members may play a role in shuttling lipids into the lipid-loading pathway of CD1 molecules. Whether these molecules play a more direct role in lipid loading into CD1 molecules, acting in a direct chaperone role, is also interesting to consider. Our data using CD1b/c chimeric tetramers, and our evidence that distinct lipids can influence the CD1-CD36 interaction, is consistent with the CD1 Ag-binding cleft being involved in this interaction. Moreover CD36L2, which appears to bind at least to CD1c, is typically retained in lysosomes, through which CD1b, c and d recycle to acquire exogenous lipid Ags^60^. It is noteworthy that CD1a, the only CD1 molecule that does not traffic to the endo-lysosome as a result of its lack of adaptor protein binding^60^, does not bind to any of the CD36 family members. Perhaps there is a lack of selective pressure to maintain CD36 binding as a result of its unique trafficking pathway. Further studies on the role of the CD36 family in CD1 Ag presentation are clearly warranted.

Our discovery that CD36 family members bind to CD1 family members has immediate implications for research into CD1-restricted T cells. Just as the advent of peptide-MHC tetramers revolutionised the study of Ag-specific T cells^2^, the application of tetramer technology to monomorphic MHC-like molecules has likewise been instrumental in the characterisation of the major innate-like T cell subsets, CD1d-restricted NKT cells^10, 11^, MR1-restricted MAIT cells^40, 61^, and Mtb-Ag-specific subsets of group 1 CD1-restricted T cells^15, 16, 17, 18^. CD1c-endo tetramers also label CD1c-restricted T cells via their TCRs^18, 33, 62^ and consecutive rounds of FACS-sort-enrichment reliably leads to the isolation of αβ and γδ T cells clones with demonstrated CD1c-TCR interaction^24^. However, prior evidence for CD1c binding to non-T lymphocytes^24^ and LILR ligands^37^ raised the question of whether CD1c tetramer staining rates overestimate the size of the CD1c-restricted T cell population if non-TCR targets can be also mediate staining. Identification of CD36 as a CD1c ligand, and blockade of this binding partner, greatly improves the sensitivity of CD1c tetramers for the identification of CD1c-restricted T cells, and indeed facilitates their use for direct ex vivo analysis of these cells. Our data suggests that CD1c-endo-tetramer^+^ T cells bind ∼0.02% of T cells, which is orders of magnitude lower than what we observed in non-anti-CD36-blocked samples. Another study that used functional measurements of T cell frequency estimated the frequency of CD1c-autoreactive T cells at ∼7% of circulating T cells^30^. In that study, large libraries of T cell clones were challenged with CD1-expressing APCs and the frequency of responding cells enumerated. The discrepancy with our tetramer-based analysis is likely a result of the different approaches. For example, the repertoire of lipids in CD1 tetramers compared to APCs may limit detection of some CD1-restricted T cells. Furthermore, in contrast to in vitro activation-based assays, CD1 tetramer staining may fail to detect low affinity TCR-binding which may be a feature of autoreactive TCRs.

There is some discrepancy in the levels of CD1c tetramer staining on non-anti-CD36-blocked samples between data presented here and that of previous studies^18, 24, 33, 62^. This will be influenced by differences in reagents, staining protocols, and instrumentation, with PE-conjugated CD1c tetramers used here being detected with a yellow-green laser to achieve maximum sensitivity. Perhaps more likely though is the pre-enrichment of T cells^33^ or overnight resting of the cells prior to staining^18, 62^ utilised in previous studies, which may be in line with the observation of reduced non-TCR-mediated staining with culture^24^. Another possible discrepancy between approaches is the PBMC isolation methods which likely result in different levels of platelet contamination. Platelets express high levels of CD36 and are also known to coat the surface of lymphoid and myeloid cells during PBMC preparation, and platelet contaminated cells can exhibit positive staining with platelet markers ^63, 64^. Because, some previous studies of CD1c-restricted T cells have isolated PBMC from platelet-pheresed^33^ or leukapheresed^18, 24^ samples these are likely to have less platelet contamination than our samples which are derived from ficoll-density gradient buffy packs. Future studies addressing the contribution of platelets to CD36-mediated CD1c tetramer staining will be informative.

In a recent study^24^, we provided evidence that some CD1c-reactive T cells recognise CD1c only when it carries lipid-antigens that have small or unobtrusive headgroups that do not interfere with binding. This may explain how CD1c-endo tetramers, which likely carry a range of lipids, can detect CD1c-autoreactive T cells because the identity of the bound lipid is not critical, only that it manages to stay out of the way of the TCR. Clearly though, CD1c is also capable of presenting specific microbial lipid-antigens such as PM and MPM to lipid-antigen-specific CD1c-restricted T cells^18^. Further, CD1c-restricted T cells appear to detect the self-lipid antigen mLPA, associated with leukemia cells^31^. It follows that CD1c-endo tetramers may only detect a fraction of the CD1c-autoreactive T cell pool in which the epitope is likely to be CD1c itself. Furthrmore, loading CD1c tetramers with specific self-lipids may allow detection of CD1c-self-reactive T cells that recognise an epitope formed from CD1c combined with a self-lipid. In line with this interpretation, our data show that CD1c loaded with specific self-lipids, such as sulfatide or GD3, can detect T cells that fail to bind CD1c-endo tetramers, suggesting that the self-lipid can contribute in a positive way to epitope formation. Thus, interpretation of all studies and enumeration of T cell precursor frequencies must consider the differing detection requirements of CD1c-specific versus CD1c-lipid-dependent autoreactivity. As we have demonstrated, the use of CD36-blockade should greatly enhance our ability to measure TCR-CD1 interactions and quantitatively study self and foreign lipid antigen-specific populations of CD1c-restricted T cells. Additionally, while non-TCR-specific staining is less problematic for other CD1 family members, we have also demonstrated how CD36-blockade can clarify CD1b and CD1d tetramer staining of fresh human T cells ex vivo. This outcome should enhance further research into all CD1b and CD1d-restricted T cell types.

Using CD1c-endo tetramers with CD36 blockade, we have been able to further characterise ‘CD1c-endo-reactive’ cells. The αβTCR repertoire of these cells was enriched for the TRBV4-1 gene, in line with previous observations^49^, but also utilised TRAV17 – a gene enriched in the CD1b-GMM-reactive TCR repertoire^51^ – as well as TRAV38-1. We also show that in some donors, a large proportion of CD1c-endo-reactive αβ T cells utilise Vδ1 and are thus ‘δ/αβ’ T cells^52^. This supports the notion that Vδ1 may have a particular predilection for CD1c^33^, as it does for CD1d^52, 53, 54, 55^. Along with the recent discovery of CD1b-reactive Vδ1^+^ γδ T cells^65^, it now appears that at least 3 members of the CD1 family can be recognised by Vδ1^+^ T cells. These observations add to a growing number of TCR genes that are similarly enriched in multiple CD1-restricted T cell populations^50^, suggesting a possible co-evolution between CD1 molecules and certain TCR genes.

Beyond TCR-repertoire, and in agreeance with a previous study^30^, CD1c-restricted αβ T cells were phenotypically diverse, including CD4, CD8 and DN subsets, as well as showing diversity in memory marker expression, suggesting differential levels of prior antigen-exposure. This is pertinent to the development of lipid-based therapeutics, as memory formation is a prerequisite for vaccine development, and diversity in co-receptor-based subsets is also preferable. Adaptive-like features also have implications for the development and regulation of these cells. The classical innate-like features of other T cells restricted to monomorphic MHC I-like molecules, such as NKT cells and MAIT cells, is thought to be acquired from their selection by CD1d- and MR1-expressing CD4^+^CD8^+^ double positive (DP) thymocytes^66^. More adaptive features are thus suggestive of a divergent developmental pathway that aligns more with conventional MHC-restricted T cells that are selected on MHC-expressing thymic epithelial cells^66^.

In summary, we have discovered a novel protein-protein interaction between CD36 family members and the CD1 family of antigen-presenting molecules. In addition to revealing an entirely new class of immunological pairing, CD36-blockade as a means to limit CD1 tetramer staining to TCR-reactivity should now facilitate further detailed studies on these T cell populations, helping to unlock key aspects of their fundamental biology, their roles in disease and homeostasis, and to aid in the development of therapeutics harnessing this knowledge. This finding furthermore paves the way for studies on the biological role of the CD36-CD1 axis.

## Materials and Methods

### Design

This study was designed to identify identify non-TCR ligands for CD1 molecules present on human blood cells.

### Human samples

Human buffy coats from healthy blood donors were obtained, with written informed consent, from the Australian Red Cross Blood Service after approval from the University of Melbourne Human Ethics Committee (1035100). Buffy coats were processed by standard density gradient using Ficoll-paque Plus (GE Healthcare) and cryopreserved in liquid nitrogen for subsequent use.

### Flow cytometry and cell sorting

All flow cytometry samples were acquired using an LSRFortessa analyser (BD Biosciences) and subsequently analysed using FlowJo v10 software (Treestar). Sorting was performed using a FACS Aria III cell sorter (BD Biosciences). For PBMC analysis, LIVE/DEAD Near-IR^+^ cells were removed, followed by doublet exclusion. Platelets were gated as FSC-A/SSC-A low, CD42b^+^ cells. Monocytes were gated as FSC-A/SSC-A high, CD14^+^ cells and lymphocytes were gated as CD45^+^ FSC-A/SSC-A Intermediate. For T cell analysis, CD14^+^ and CD19^+^ cells were removed from analysis.

### Ex vivo staining of human PBMCs

PBMCs were incubated in DNAse for 30 min at 37°C prior to being filtered. For experiments examining CD1c-restricted T cells, cells were washed in PBS and incubated in PBS + 50 nM dasatinib (Stem Cell Technologies, Canada) + DNAse at 37°C for 30 min. Cells were then blocked with human Fc-block (Miltenyi) in the presence or absence of purified anti-CD36 antibody (Clone 5-271; Biolegend) at 10 μg/ml for a further 15 min at 4°C. CD1c tetramers were then added directly to blocking cocktail at a final concentration of 1 μg/ml and stained for 30 min at room temp. Cells were washed once before being stained with surface mAb (Table S1) and viability dye LIVE/DEAD Fixable Near-IR (Thermo-Fisher Scientific, USA) for 30 min at 4°C. Cells were then washed twice, fixed in 2 % PFA, washed once more and filtered prior to immediate FACS acquisition. For dual tetramer labelling experiments, cells were stained with mAb cocktail containing the first tetramer, followed by avidin and biotin blocking (Dako) before staining with the second tetramer. For staining of δ/αβ T cells, enriched cells were stained with anti-Vδ1 clone A13 supernatant (a gift from L. Moretta) for 30 min at 4°C, washed thrice, then stained with anti-mouse IgG, washed three times, blocked with 5 % normal mouse serum, and then stained with other surface mAb.

### Magnetic enrichment

Human PBMCs were isolated as above. Cells were then treated with dasatinib for 1 hr at 37°C prior to blocking with Fc-block (BD Bioscience), 5 % normal mouse serum, and purified anti-CD36 for 20 min at RT. Cells were then stained with CD1c-endo tetramer-PE for 30 min at RT, washed twice and then incubated with anti-PE magnetic microbeads (Miltenyi) for 20 min at 4°C. Cells were washed twice more, followed by magnetic enrichment using LS MACS enrichment columns (Miltenyi). Enriched cells were then stained with surface mAb as above and subsequently FACS-sort purified or FACS analysed.

### Staining of cell lines

Cell lines were maintained in RF10 complete media consisting of RPMI-1640 base supplemented with 10 % FBS (JRH Biosciences), Penicillin (100 U/mL), Streptomycin (100 μg/mL), Glutamax (2 mM), sodium pyruvate (1 mM), nonessential amino acids (0.1 mM), HEPES buffer (15 mM), pH7.2-7.5 (all from Invitrogen, Life Technologies) and 2-mercaptoethanol (50 μM, Sigma). Cells were harvested, washed with PBS and stained for 30 min at RT with antibody cocktails including LIVE/DEAD Near-IR, tetramers and/or mAbs (see table 1). Cells were then washed and filtered through 100 μm mesh prior to immediate acquisition by flow cytometry.

### In vitro T cell expansion

Sorted cells were stimulated with platebound anti-CD3 (10 μg/ml; clone OKT3; BD Biosciences) and anti-CD28 (2 μg/ml; clone CD28.2; BD Bioscience) in the presence of rhuIL-2 (200 U/ml; Peprotech), rhuIL-7 (50 ng/ml; eBiosciences) and rhuIL-15 (5ng/ml; eBiosciences). After 48 hrs, cells were removed from stimulus and allowed to expand for approximately 10-14 d in cytokine media. Cells were assessed directly or cryopreserved for subsequent analysis. T cell culture media consisted of a base of 1:1 RPMI-1640 and AIM-V media (Invitrogen, Life Technologies), supplemented as per RF10 above with the addition of 2 % human AB serum (Sigma).

### Single cell TCR sequencing

For sequencing of TCRs in Table 1, magnetically enriched CD1c-endo tetramer^+^ cells were FACS-purified and expanded in vitro as above prior to restaining with CD1c-endo tetramers and scSorting. Human T cell receptor genes were then sequenced as previously described^67^. In brief, single CD1c tetramer^+^ T cells were sorted into 96-well PCR plates. cDNA was generated using SuperScript VILO cDNA synthesis kit (ThermoFisher). cDNA was then subjected to 2 rounds of semi-nested PCR using multiplexed primer sets to specifically amplify the TCR-α and TCR-β chains. Amplified DNA was sequenced using Sanger Sequencing, and sequences were analysed using the IMGT/V-QUEST online tool. For sequencing of TCRs in figure 5, cells were single cell sorted directly ex vivo and PCRs performed as above.

### Tetramers

CD1 tetramers. Human CD1a, CD1b, CD1c and CD1d tetramers were produced as previously described^22, 41^. In brief, 293S.GnTI^-^ cells were co-transfected using PEI transfection reagent with pHLSec expression vectors encoding the truncated ectodomain of CD1b, CD1 or CD1d together with a separate pHLSec vector encoding full length human β2-microglobulin (β2m). For CD1a, a single vector encoding the truncated ectodomain of CD1a linked to β2m via carboxy-terminal leucine zippers was used. For chimeric CD1c/b proteins, constructs were based on a previous publication^44^ which linked β2m to the truncated CD1 ectodomain via a glycine-serine linker (GGGGSGGSGSGGGSS) and fused the CD1c α1 and α2 domains (A1-V186) to the CD1b α3 domain (K186-P280), followed by C-terminal AVI-tag and 6xHIS tag. Recombinant CD1 proteins were purified using Ni-NTA agarose, biotinylated with BirA enzyme and further purified by size-exclusion chromatography. MR1-5-OP-RU tetramers were produced as previously described^68^. In brief, truncated ectodomains of human MR1.C262S were expressed as inclusion bodies in Escherichia coli (strain BL21) along with human β2-microglobulin (β2m). MR1 and β2m inclusion bodies were then refolded in vitro in the presence of 5-A-RU (Fairlie Laboratory, University of Queensland) and methylglyoxal (Sigma) using oxidative refolding, prior to dialysis and subsequent purification using Ni-NTA agarose (ThermoFisher). Human MR1-5-OP-RU were enzymatically biotinylated using BirA enzyme (produced in-house) followed by further purification by size exclusion chromatography prior to storage at −80°C. All monomers were tetramerised using streptavidin-PE, -PE-Cy7 or -BV421 (all BD Pharmingen).

### Lipids

Lipids were dissolved by sonication in TBS (10 mM Tris pH 8.0, 150 mM NaCl) containing 0.05 % (v/v) tyloxapol (Sigma) and stored at −20°C prior to use. Prior to loading, lipids were thawed and re-sonicated for 30 min. All loading was performed overnight at RT. For staining in figure 1, CD1d was loaded with α-GalCer (KRN7000) at a 3:1 molar ratio (lipid:CD1). For Figure 6, CD1d was loaded with PBS44 (provided by Paul B. Savage, Brigham Young University) at 6:1, LPC (C18:1) or sulfatide (C24:1), both at 12:1. CD1c was loaded with LPC (C18:1), sulfatide (C24:1), PC (C18:0-C18-1) or GD3 at 12:1. KRN7000, LPC, PC, sulfatide and GD3 were purchased from Avanti Polar Lipids. CD1b was loaded with GMM (provided by D. Branch Moody, Brigham and Women’s Hospital, Boston) at 6:1.

### Transient transfections

Full-length CD36, CD36L1 and CD36L2 genes, as well as TCRα- and β-chain genes linked by a p2A-linker were cloned the pMIGII expression vector. Vectors were used to transiently transfect 293T or 293T.CD36L1^-/-^ cells using FUGENE (Promega). TCR transductions also included a pMIG vector encoding CD3ε, δ, γ and ζ, Cells. Cells were harvested after 48 hr, washed once in PBS + 2% FBS, then stained for 30 mins at RT with CD1 tetramers. Cells were then washed twice, fixed in 2% PFA for 15 min at RT before a final wash step.

### Jurkat cell lines

Stable TCR-expressing cell lines were generated by retroviral transduction of parental Jurkat76 cells using pMIGII plasmids above, as previously described^54, 69^.

### Generation of CRISPR/Cas9-mediated knockout cell lines

For CD36L1 knockout lines, lentiCRISPRv2GFP plasmid^70^ was a gift from David Feldser (Addgene plasmid #82416; http://n2t.net/addgene:82416; RRID:Addgene_82416) and lentiCRISPRv2-mCherry was a gift from Agata Smogorzewska (Addgene plasmid #99154; http://n2t.net/addgene:99154; RRID:Addgene_99154). sgRNA cloning was performed as described by the GeCKO Lentiviral CRISPR toolbox (Zheng Lab, Broad Institute, MIT), using the following oligos designed using the crispr.mit.edu tool: 1. CACCGTCATGAAGGCACGTTCGCCG and 2. CAGTACTTCCGTGCAAGCGGCCAAA to generate an sgRNA specific for the target sequence CGGCGAACGTGCCTTCATGA within the CD36L1 gene. 293T cells were then transiently transfected with both GFP and mCherry plasmids using Fugene HD reagent. After 3 days of transfection, cells were harvested and single cell sorted for high GFP and mCherry co-expression plus low CD36L1 expression. After in vitro propagation for 2-3 weeks, clones were screened for CD36L1 expression compared to wildtype 293T cells. One 293T.CD36L1-KO cell line was further propagated, and aliquots cryopreserved for subsequent experiments.

### Whole genome CRISPR/Cas9 knockout screen

CRISPR library screening was performed as previously described^71^ following the Zheng lab protocol (Broad institute, MIT). In brief, the human GeCKO v1 sgRNA library was lentivirally transduced into C1R cells in 3 biological replicate libraries as previously described^45^. Each of the 3 libraries were split into two, and half stained with CD1c tetramers and tetramer-negative cells enriched using FACS. The other half of each library was maintained in culture unsorted. Enrichment was performed twice to enrich for C1R cells containing sgRNAs that result in loss of tetramer staining. Genomic DNA was then extracted from FACS-enriched samples and unsorted libraries. The sgRNAs in each DNA sample were amplified using the two-step PCR method described previously^71^ using Hot Start Taq Polymerase (New England Biolabs), however in the second PCR the 8bp barcode was incorporated into the reverse as well as forward primer. Barcoded amplicons were pooled and purified using Nucleo Mag NGS beads (Macherey-Nagel), then sequenced using HiSeq 2500 (Illumina). Sequencing reads were matched to known sample-specific barcodes and gene-specific sgRNA sequences to obtain a count matrix. sgRNAs were retained if they had a non-zero count in at least 3 samples and a total count greater than 8 across all samples. The filtered count matrix was subject to TMM normalization^72^ using *edgeR* v3.20.9^73^. Generalized linear models were fitted to the counts from each sgRNA to look for differential abundance between the replicate sorted and unsorted samples using likelihood ratio tests^74, 75^, with sgRNAs ranked according to their false discovery rate (FDR)^76^. A volcano plot of the −log_10_(*p*-value) versus log_2_-fold-change (logFC) was generated, and sgRNAs with FDR < 0.05 and logFC > 0 were highlighted in interactive plots using *Glimma* v1.6.0^77^.

### Recombinant monoclonal antibody production

DNA constructs encoding anti-SCARB2 antibody heavy and light chain variable domains (clone JL2^78^) were synthesized (ThermoFisher) and cloned into mammalian expression vectors containing a mouse IGHV signal peptide and IgG1 constant regions. Antibodies were expressed by transient transfection of mammalian Expi293F cells and purified using Protein G column chromatography (GE), followed by buffer-exchange into PBS (**Fig. S6**).

### Statistical Analysis

Statistics were generated using Graphpad Prism software. Details of individual experiments are included in associated figure legends.

## Data availability

The authors declare that the data supporting the findings of this study are available within the article, supplementary information files and source data, or are available upon reasonable requests to the authors.

---

**Figure S1:**
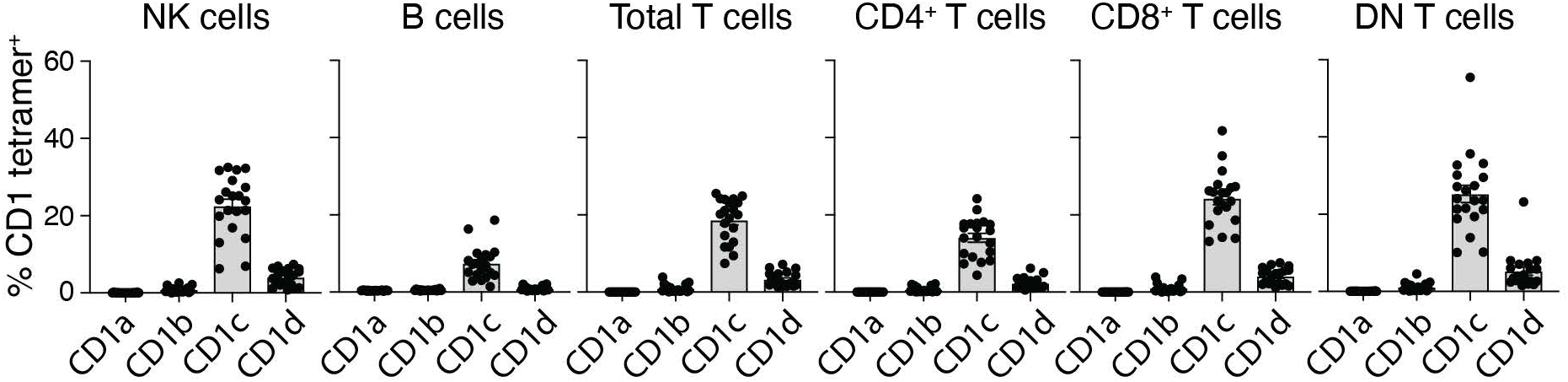
CD1-endo tetramer staining of lymphoid subsets. Bar graphs showing the proportion of cells within different lymphoid subsets of PBMC staining with each CD1-endo tetramer (n=20). DN = CD4-CD8-double negative. NK cells defined as CD45+CD3-CD56+, B cells defined as CD45+CD3-CD19+, T cells defined as CD45+CD3+CD19-.

**Figure S2:**
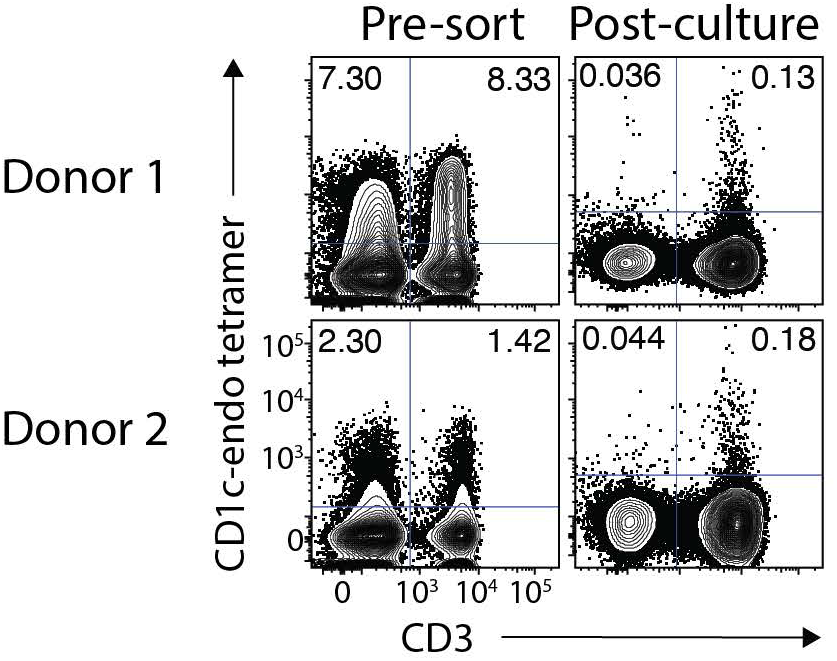
In vitro culture reduces non-TCR-mediated CD1c-endo tetramer staining on human T cells. Contour plots showing CD3+ T cells from PBMC at the time of sorting (left) and after 1O days of in vitro expansion (right) on 2 example donors.

**Figure S3:**
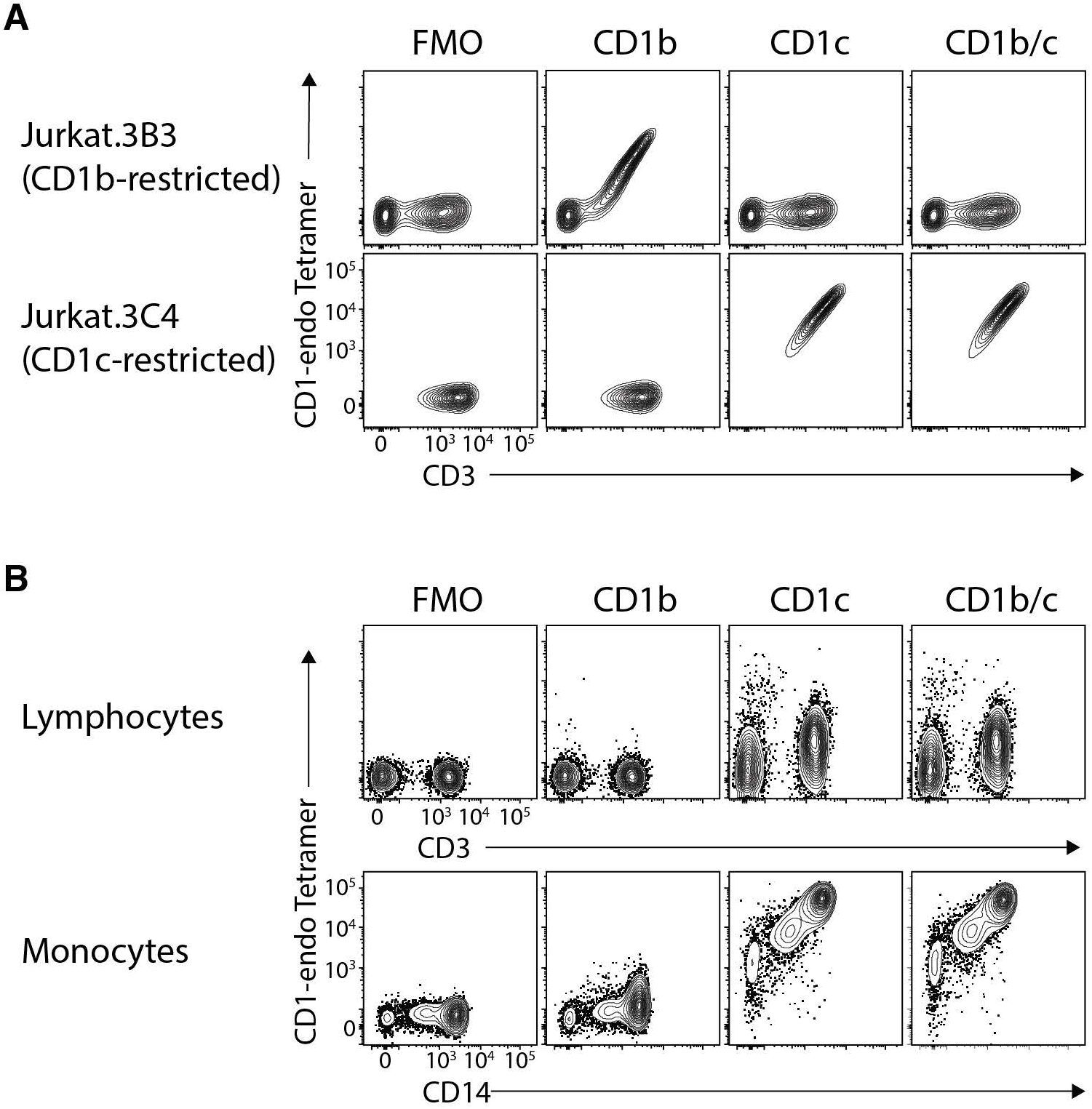
CD1b/c chimeric tetramers stain CD1c-restricted T cells, but exhibit non-TCR mediated PBMC staining. **A.** Contour plots showing CD1 tetramer staining on jurkat cells transduced to express CD1b-restricted TCR 383 or CD1c-restricted TCR 3C4. **B.** Contour plots showing CD1 tetramer staining on lymphocytes and monocytes. CD1b/c = chimeric CD1b/c protein.

**Figure S4:**
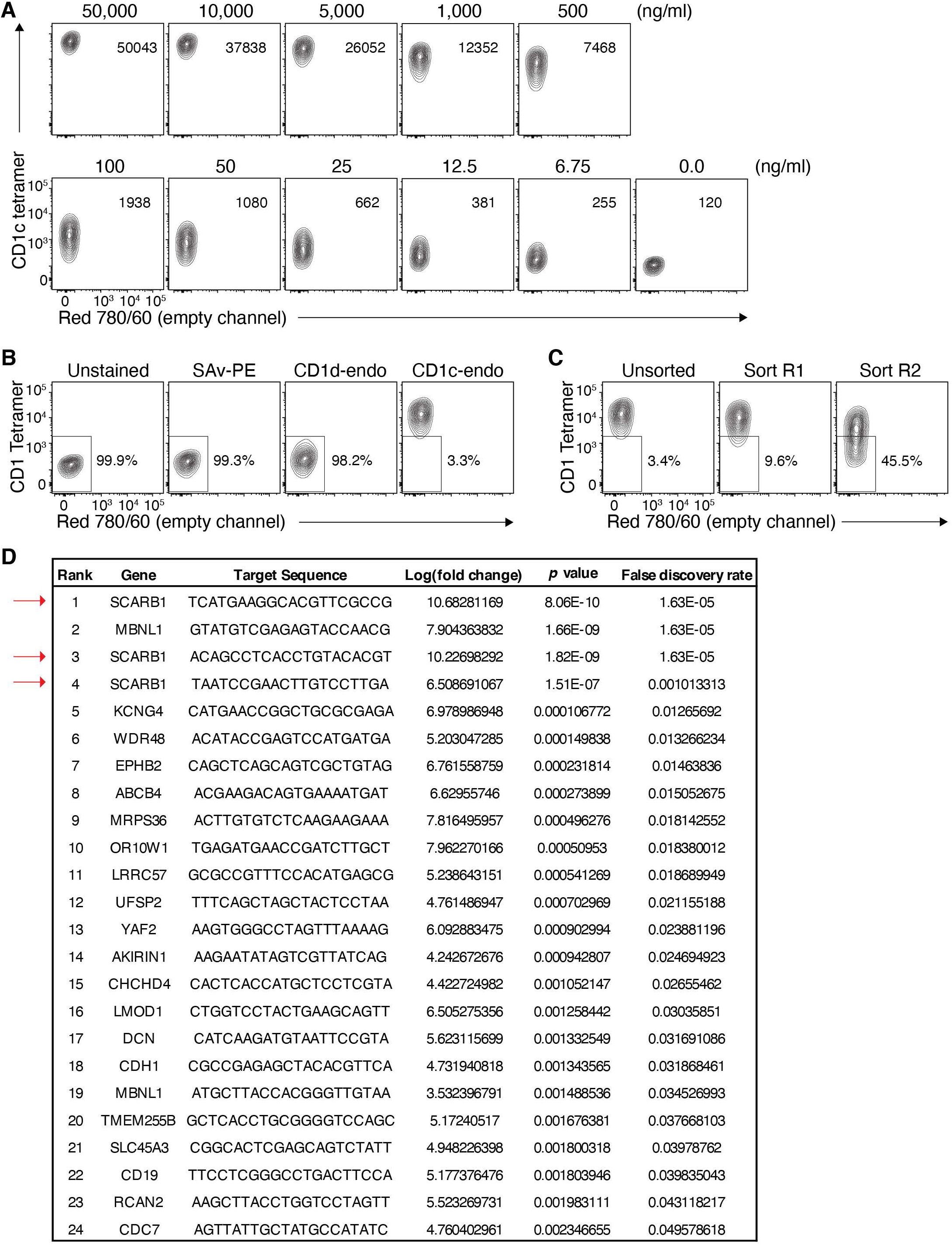
CRISPR GeCKO screen supplementary data. **A.** Contour plots showing a dose titration of CD1c-endo tetramer staining on wildtype C1R cells, establishing 1µg/ml as the minimum staining dose required to maintain staining of all cells. **B.** Contour plots showing CD1 tetramer staining at the 1µg/ml dose, of C1R cells transduced with the GeCKO v1 human CRISPR KO library. **C.** Contour plots showing CD1c-endo tetramer staining of the unsorted library compared to the library after 1 or 2 rounds of enrichment for CD1c-endo tetramer-negative cells. **D.** Table showing all significantly upregulated gRNA hits with a false-discovery rate <0.05 in the CD1c-tetramer^low^ population compared to control unsorted cells. Red arrows highlight SCARB1 (CD36L1) guides.

**Figure S5:**
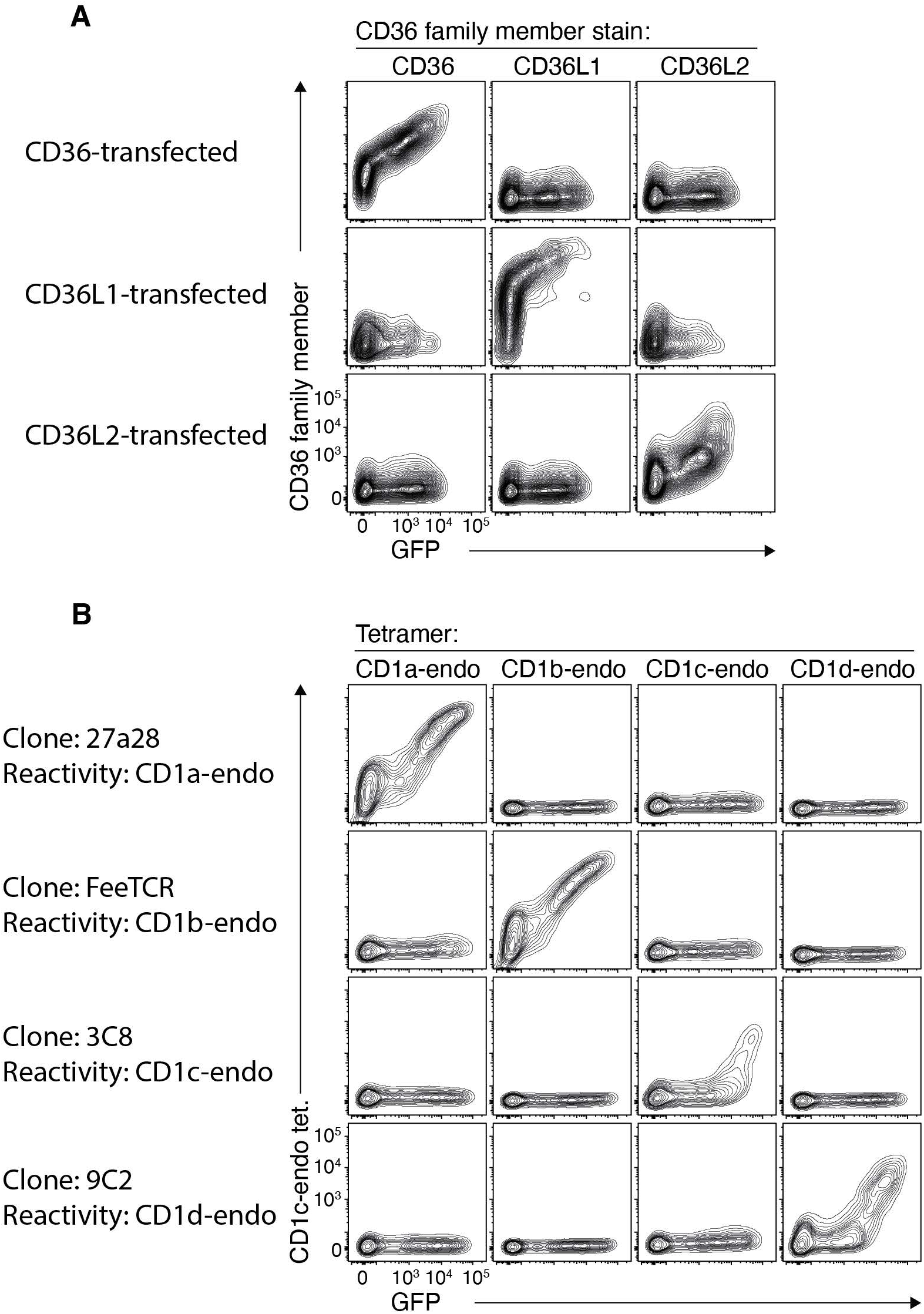
Controls for 293T transient transfections with CD36 family members. **A.** Contour plots showing staining for CD36 family members on 293T.SCARB1^-/-^ cells transiently transfected to express CD36, CD36L1 or CD36L2. **B.** Contour plots showing CD1 tetramer staining of 293T.SCARB1^-/-^ cells transiently transfected to express CD1-restricted TCRs. Data presented in (A) and (B) are representative of 3 independent experiments each.

**Figure S6:**
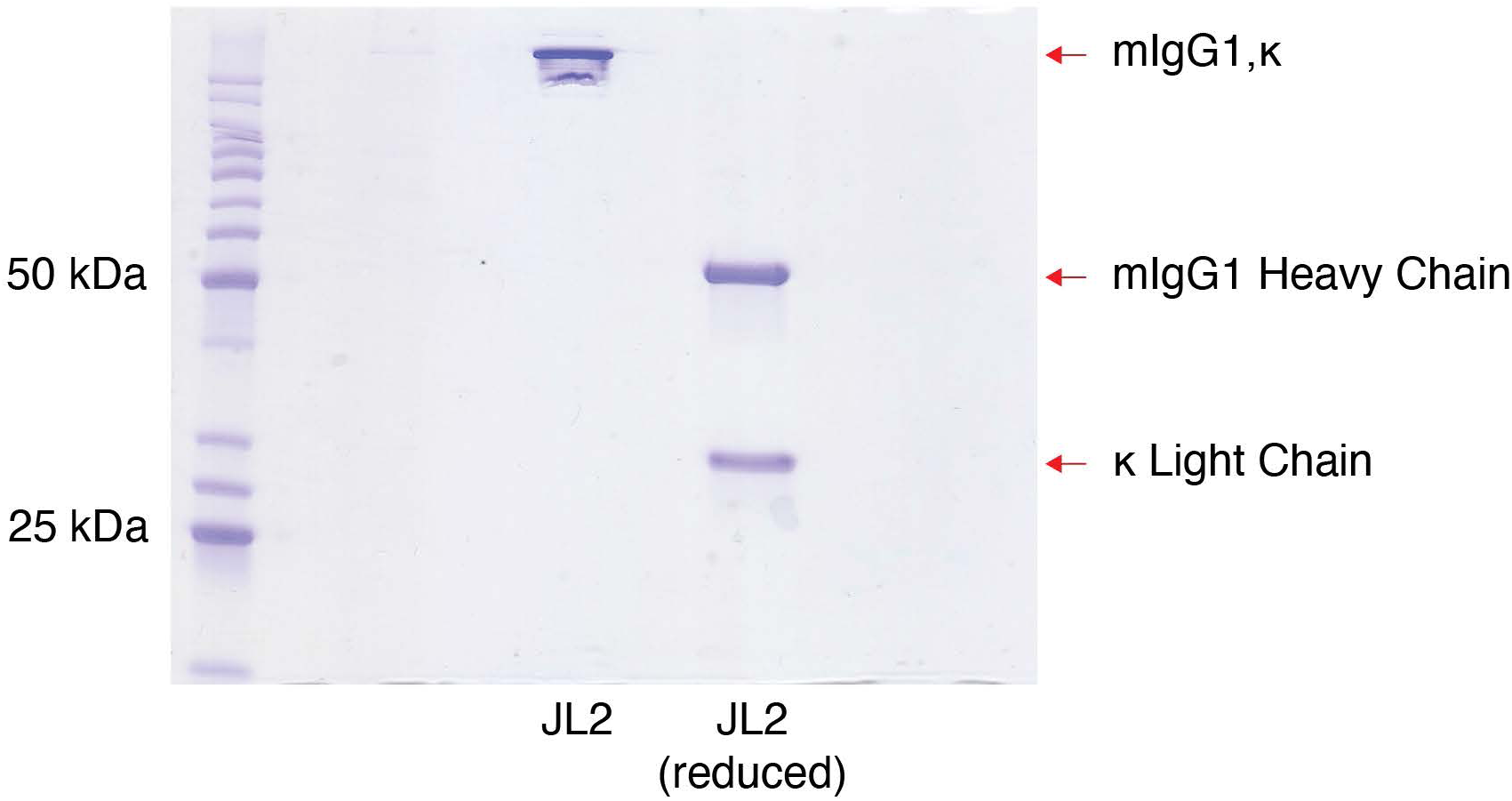
JL2 antibody production. SOS-PAGE gel of purified recombinant JL2 monoclonal antibody under reduced and non-reduced conditions.

**Table S1.** List of monoclonal antibodies

| Epitope | Clone | Fluorophore | Dilution | Vendor |
| --- | --- | --- | --- | --- |
| human CD3e | UCHT1 | BUV395 | 1:50 | BD |
| human CD3e | UCHT1 | BV421 | 1:50 | BD |
| human CD4 | SK3 | BUV496 | 1:50 | BD |
| human CD4 | RPA-T4 | AF700 | 1:100 | BD |
| human CD8a | RPA-T8 | BV650 | 1:50 | BD |
| human CD14 | M5E2 | BUV805 | 1:300 | BD |
| human CD14 | MoP9 | APC-Cy7 | 1:50 | BD |
| human CD19 | 2J25C1 | APC-Cy7 | 1:50 | BD |
| human CD19 | H1B19 | PE-Dazzle | 1:100 | Biolegend |
| human CD27 | O323 | BV786 | 1:50 | BD |
| human CD36 | 5-271 | APC | 1:50 | Biolegend |
| human CD36L1 | m1B9 | APC | 1:50 | Biolegend |
| human CD42b | HIP1 | APC | 1:50 | Biolegend |
| human CD45 | 2D1 | AF700 | 1:50 | Biolegend |
| human CD45RA | HI100 | PerCP-Cy5.5 | 1:100 | BD |
| human CD56 | HCD56 | BV605 | 1:100 | Biolegend |
| human TRAV1-2 | 3C10 | APC | 1:50 | Biolegend |
| human TCRgd | 11F2 | FITC | 1:50 | BD |
| mouse IgG1 | Poly4053 | BV421 | 1:40 | Biolegend |

## Acknowledgements

We are grateful to Dr. Paul Savage (Brigham Young University, UT, USA) for providing α-galactosylceramide analogue PBS44 used for production of CD1d-α-GalCer tetramers, Prof. David Fairlie and Jeffrey Mak (University of Queensland, Australia) for providing 5-A-RU used for production of MR1-5-OP-RU tetramers, Prof. Lorenzo Moretta (Bambino Gesu Children’s Hospital, Rome, Italy) for anti-Vδ1 clone A13 supernatant and Dr. Adam Wheatley (Peter Doherty Institute, University of Melbourne, Australia) for plasmids used for JL2 mAb production. We thank staff from the flow cytometry facilities at the Department of Microbiology and Immunology at the Peter Doherty Institute and the Melbourne Brain Centre at the University of Melbourne. This work was supported by the Australian Research Council (ARC; CE140100011) and the National Health and Medical Research Council, Australia (NHMRC; 1113293 and 1145373) and the National Institutes of Health, USA (NIH; R01 AR048632). DIG was supported by an NHMRC Senior Principal Research Fellowship (1117766) and DGP is supported by a CSL centenary fellowship.

## Author contributions

N.A.G. performed experiments and/or directed the experimental work. S.J.R., C.F.A, K.H.A.G., R.S., R.D.R., C.V.N-R., D.B.M., D.G.P and A.P.U. performed or assisted with cellular experiments. H.E.G.W., S.L., and J.A.V. assisted with CRISPR experiments and S.S. and M.E.R. performed CRISPR screen data analysis. N.A.G. and D.I.G. conceived the study, co-wrote the paper and contributed equally.

